# Organoid-T cell co-cultures functionally stratify tumor-reactive T cells and their responses to immune checkpoint inhibitors

**DOI:** 10.64898/2026.05.27.727968

**Authors:** Elliot Merritt, Julie-ann Cavallo-fleming, Genesis Lara Granados, Shalini Nath, Wooseung Lee, Ramja Sritharan, Subhasree Sridhar, Laura Zuluaga, Edgardo V. Ariztia, Fred R. Hirsch, Martin Walsh, John P. Sfakianos, Ketan Badani, Rachel Brody, Amir Horowitz, Alexander M. Tsankov, Daniela Sia, Benjamin Hopkins, Anna S. Tocheva

**Author notes:** Corresponding authors: Anna S. Tocheva, PhD, Benjamin Hopkins, PhD. These authors contributed equally (Benjamin Hopkins, Anna S. Tocheva).

## Abstract

Tumor-reactive T cells (TRTs) are critical for anti-tumor immunity but are incompletely captured by current assays, which fail to reproduce tumor-specific antigen diversity. Here, we show that multiplex functional profiling of patient-derived tumor organoid-T cell co-cultures (PDOTs) enables robust identification of TRTs across CD8, CD4, and double-negative (DN) T cell populations. Single activation markers underestimated TRT responses, whereas integrated analysis revealed broader functional repertoire. MHCI blockade abrogated CD8 and DN TRT responses while preserving CD4 reactivity, supporting antigen-dependent recognition across T cell lineages. Tumor PDO expressed MHCI and MHCII, and PDOTs enabled generation and detection of TRTs from peripheral blood. PD1 blockade induced heterogeneous responses, enhancing CD8 and DN activity and unexpectedly augmenting CD4 reactivity. PDOTs further identified additional inhibitory pathways whose therapeutic targeting in combination with PD1 blockade increased TRT responses. These findings establish PDOTs as a platform to identify TRTs and functionally stratify patient-specific tumor-T cell responses to checkpoint immunotherapy.

## Introduction

Tumor-derived 2D lines and animal models do not represent the genetic heterogeneity of patient tumors limiting their ability to recapitulate tumor-specific antigens and oncogenic programs that shape immune responses and tumor-T cell interactions (1,2). These shortcomings highlight the need for patient-derived models that preserve the malignant programs and antigen diversity of primary tumors, enabling the accurate measurement of tumor-reactive T cell responses.

Patient-derived tumor organoids (PDOs) have emerged as a powerful platform to address these challenges by preserving the genotype, antigen repertoire, and cellular programs of primary tumors across diverse cancer types^1-4^. Unlike 2D cell lines, which undergo genetic drift and gradually become monoclonal, PDOs retain the tumor heterogeneity and three-dimensional architecture of primary tumors^5,6^. Comparative studies have demonstrated that tumor PDOs recapitulate patient responses to chemotherapy^7^ and chemoradiation^8^, and identify effective strategies to overcome therapeutic resistance^8,9^. As such, PDOs provide a physiologically relevant and scalable platform to study human cancer biology and evaluate therapeutic strategies that are infeasible in human subjects.

T cells are central mediators of anti-tumor immunity and key targets of modern cancer treatments, including immune checkpoint inhibitors (ICIs), bispecific antibodies and adoptive T cell approaches. As these therapies become standard of care, there is a growing need for preclinical platforms that capture the functional complexity of anti-tumor T cell interactions. Organoid co-cultures with autologous blood- or tumor-derived T cells (PDOTs) have been used to study tumor-T cell interactions^4,10,11^, enrich tumor-reactive T cells^3^, and evaluate T cell responses to PD1 ICIs^12^. However, these studies have largely focused on CD8 T cell reactivity and relied on limited functional readouts restricting the detection of the full tumor-reactive T cell repertoire and functional heterogeneity of ICI responses.

Here, we show that PDOT co-cultures combined with multiple activation-induced marker (AIM) profiling enable the comprehensive identification of tumor-reactive T cells (TRTs) across CD8, CD4 and double negative (DN) T cells (**Fig. 1A**). We demonstrate that PDOs can be leveraged to identify and enrich TRTs from both tumor infiltrating lymphocytes and peripheral blood. Using this platform, we functionally characterize responses to PD1 blockade and identify candidate combination ICI strategies that enhance anti-tumor T cell responses. Together, this work establishes PDOT co-cultures as a robust and adaptable platform for functional interrogation of tumor-reactive T cells and personalized evaluation of patient-specific sensitivity to immunotherapy.

**Figure 1.**
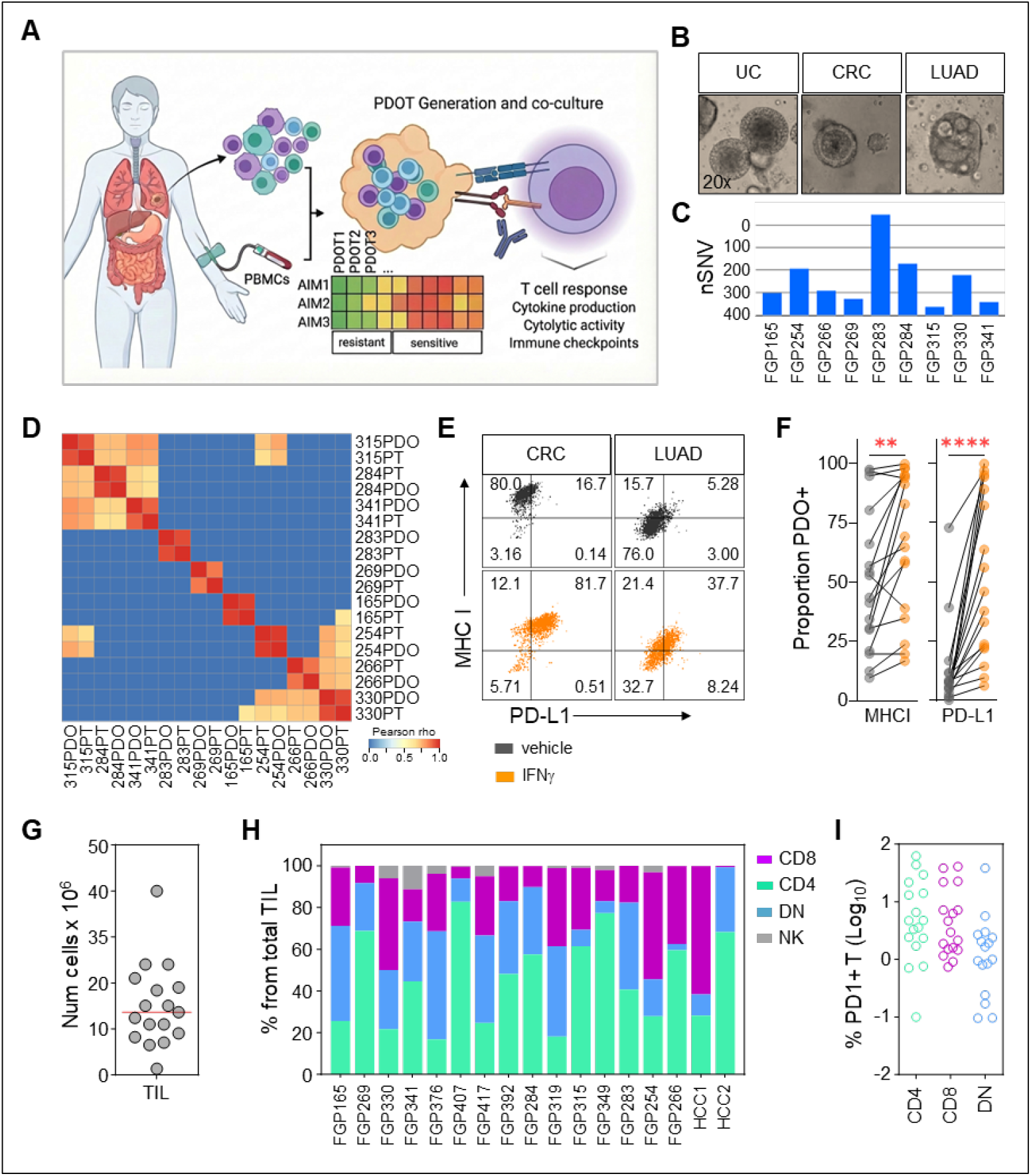
Study design and generation of tumor PDO and autologous TIL. **A**, Graphical summary. PDOT are generated from autologous PDO and T cell derived from TIL or peripheral blood. Multiplex functional profiling of PDOT co-cultures evaluates tumor T cell reactivity and responses to immune checkpoint inhibitors. **B**, Example brightfield micrographs of tumor PDO. **C**, Number of nSNV detected per PDO assayed using whole exome sequencing. **D**, Somatic SNV concordance between tumor PDO and primary patient tumors as quantified using Pearson correlation coefficient (color bar) of variant allele frequencies between samples and visualized using a heatmap following hierarchical clustering (Ward linkage). **E**, Representative flow cytometry plots of MHCI and PD-L1 expression in PDO following vehicle (grey) or IFN_γ_ (orange) stimulation. **F**, Proportion of MHCI+ or PD-L1+ cells in tumor PDO (n=17) following IFN_γ_ stimulation; **p<0.01, ****p<0.0001 using paired Student’s t test **G**, Number of total TIL generated from each patient’s tumor dissociate (n=17). **H**, Proportion of NK and CD4, CD8 and DN T cells in generated TIL lines. **I**, Proportion of PD1+ TIL across CD4, CD8 and DN T cells.

## Results

### Generation of tumor PDO and paired tumor-derived T cells

We generated a panel of 17 patient-derived tumor organoids (PDO) with paired tumor-infiltrating lymphocytes (TIL) from resections or core needle biopsies of liver, colon, lung, testicular and kidney cancers using our previously described pipeline^13,14^ (**Fig. 1A-B, Supplementary Fig. S1**). We performed whole exome sequencing for nine of the PDO samples and paired primary tumors to confirm genomic concordance. The PDO had variable tumor mutational burden evaluated by the number of non-synonymous single nucleotide variants (nSNVs) (**Fig. 1C**) and showed high genomic concordance with their matched primary tumors (**Fig. 1D)**. Since MHC class I (MHCI) expression is key to CD8 T cell mediated anti-tumor immunity, while tumor PD-L1 interferes with T cell responses and is an immunotherapeutic target, we determined MHCI and PD-L1 expression in the PDO in response to 18-hour IFN-gamma (IFN_γ_) stimulation as described previously^4,15^ (**Fig. 1E**). The majority of PDO (14/17 PDO) expressed high MHCI levels prior to stimulation, which was further enhanced in response to IFN_γ_ (**Fig. 1F**). As expected, PD-L1 levels were low prior to stimulation and IFN_γ_ increased PD-L1 expression in all tumor PDO (**Fig. 1F and Supplementary Fig. S1**). In parallel, we generated autologous in vitro TIL lines from each tumor dissociate (**Fig. 1G**). Most of the cells in TIL cultures were T cells (>85%) (**Fig. 1H and Supplementary Fig. S2A**) with good representation of PD1+ CD4, CD8 and DN populations (**Fig. 1I and Supplementary Fig. S2B**). Collectively, we show that PDO preserved the genomic features of primary patient tumors and can be reliably generated in parallel with autologous tumor-derived T cells.

### Multiplexed functional analysis improves detection of tumor-reactive T cells in PDOT co-cultures and reveals CD4 and DN-mediated reactivity

We next determined whether PDOT co-cultures can identify TRTs from expanded TIL lines. We hypothesized that conventional assays relying on limited functional markers underestimate the breadth of the functional TRT repertoire. To address this, we designed a multiplex activation-induced marker (AIM) panel capturing cytokine production and cytolytic activity (**Supplementary Fig. S3A**). Polyclonal stimulation of the TIL lines with PMA/Ionomycin confirmed that individual T cells exhibited distinct and non-overlapping functional profiles, indicating that single markers are insufficient to capture the full TRT repertoire (**Supplementary Fig. S3B-C**). Following, we pre-stimulated the PDO with IFN_γ_ to enhance antigen presentation and co-cultured tumor-organoids with autologous TILs to determine the proportion of total activated T cells in the co-cultures for each marker relative to unstimulated T cells (**Fig. 2A**). The number of AIM+ T cells in the PDOTs varied between 0.01%-15% (**Fig. 2B**) and we detected T cell reactivity in 10/17 PDOTs (**Fig. 2C**). However, consistent with our polyclonal stimulation results, individual markers failed to capture the entirety of TRT responses across PDOTs (**Supplementary Fig. S3C)**. For example, no IFN_γ_ was detected in the PDOTs in **Fig. 2D**, while GM-CSF, TNFα and CD137 were produced. Similarly, in **Supplementary Fig. S3C** the proportion of IFN_γ_-producing or degranulating TRT underrepresents the total T cell reactivity in the co-cultures captured by the addition of GM-CSF and TNFα. To overcome this limitation, we applied Boolean integration of all AIMs to quantify total TRT responses. This approach significantly increased the detection of TRT across PDOTs (**Supplementary Fig. S3D**) and revealed that seven PDOTs had TRT responses above the unstimulated background control (**Fig. 2E**). To confirm antigen-specific reactivity, we blocked MHCI presentation (**Fig. 2A**, bottom panel). While MHCI blockade reduced CD8 and DN TRT responses (**Fig. 2F-G** and **Supplementary Fig. S3E-F**), it was unable to decrease TRT responses in all PDOT cultures. Of note, DN T cells were not CD8 T cells that down-modulated the CD8 co-receptor since we performed extracellular and intracellular anti-CD8 antibody staining to identify true DN T cells. Instead, CD4-mediated reactivity was preserved (**Fig. 2G**), indicating that PDOT co-cultures capture both MHCI-restricted and non-MHCI-restricted T cell responses. Subsequent analysis revealed that TRTs were distributed across CD8, CD4 and DN T cells (**Fig. 2H**). Notably, a substantial fraction of TRT responses was mediated by CD4 and DN T cells. Consistent with this, tumor PDO expressed MHCII, which was further enhanced by IFN_γ_ stimulation (**Fig. 2I**), supporting the presence of CD4-mediated tumor recognition. Collectively, these data show that PDOT co-cultures combined with multiplex functional profiling enable robust detection of tumor-reactive T cells across CD8, CD4 and DN lineages.

**Figure 2.**
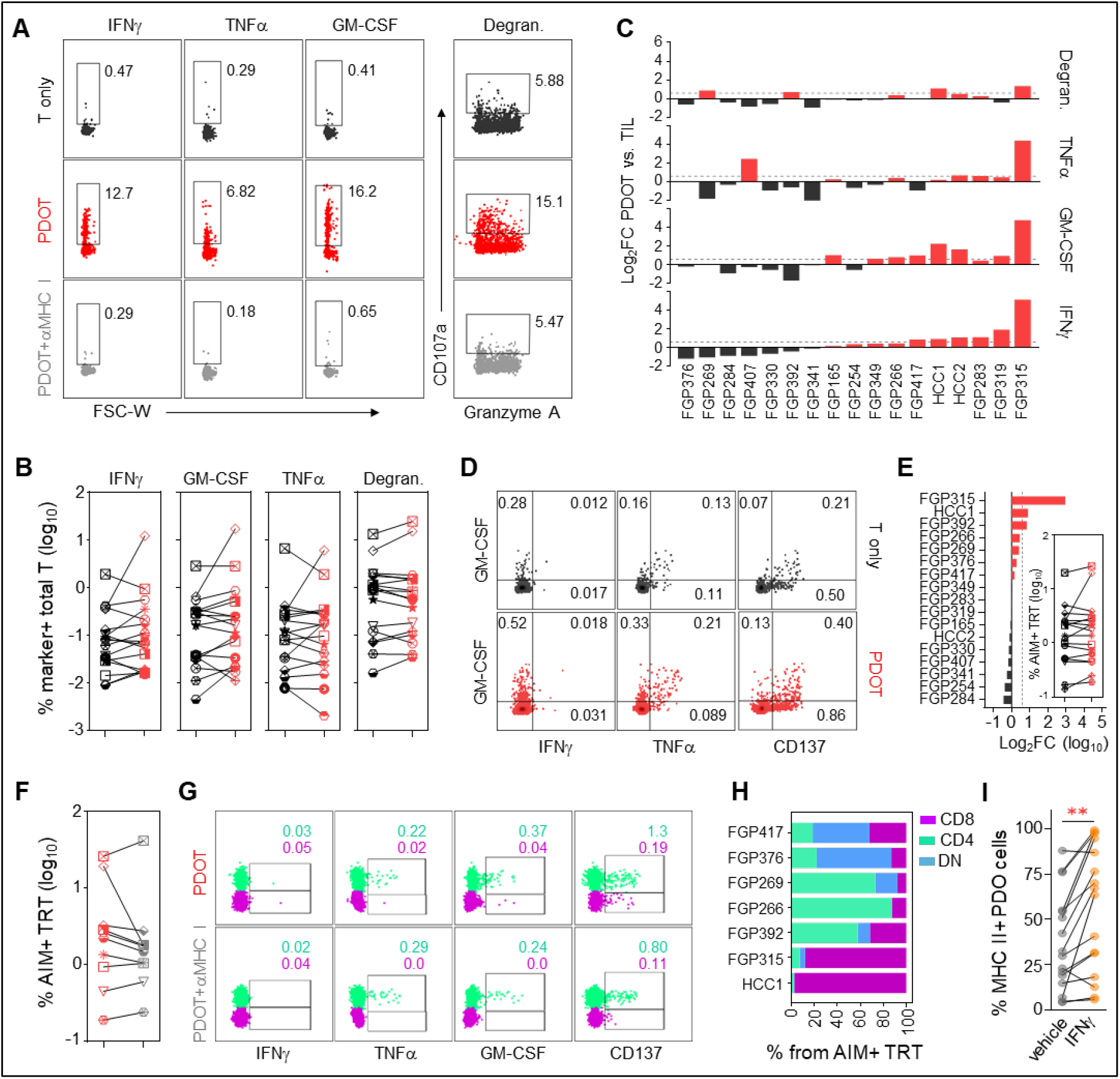
PDOT co-cultures identify CD8, DN and CD4 tumor-reactive T cells. **A**, Representative flow cytometry plots from PDOT co-culture showing intracellular cytokine staining of unstimulated TIL (black) or gated on total T cells in the PDOT co-culture (red) and PDOT in the presence of anti-MHCI blocking antibody (light grey). **B**, Proportion of marker positive unstimulated T cells alone (black) or in PDOT (red) (n=17). **C**, Log_2_FC of total TRT proportion in PDOT relative to T cells alone. **D**, An example of TRTs in the PDOTs that produce TNFα, GM-CSF and CD137 but not IFN_γ_ in response to autologous PDO. **E**, Log_2_FC responses of total TRT identified by Boolean gating of all upregulated functional markers relative to T cells alone. Inset, proportion of AIM+ TRT estimated by Boolean integration. **F**, Boolean TRT response measured across all AIMs in PDOTs (red) and in PDOTs cultured in the presence of anti-MHCI blocking antibody (light grey). **G**, Representative dot plots of CD4 and CD8 responses in FGP266 showing that MHCI blockade does not reduce CD4 T cell reactivity in the PDOT. **H**, Distribution of CD4, CD8 and DN T cells across TRTs in each baseline reactive PDOT from E. **I**, Proportion of MHCII+ cells in PDO in response to vehicle or IFN_γ_; **p<0.01 using paired Student’s t test.

### PDOT co-cultures enable antigen-agnostic generation of tumor-reactive T cells from peripheral blood revealing a circulating TRT repertoire that spans CD8, DN and CD4 T cells

Next, we asked whether TRTs could be generated from peripheral blood using PDOT co-culture enabling access to circulating TRT populations. Optimized protocols for the efficient expansion and enrichment of functional TRTs from peripheral blood are needed for the identification of the circulating TRT repertoire, isolation of tumor-specific T cell receptors and culture of non-exhausted TRTs for downstream adoptive therapies. Therefore, we determined whether PDO can be used for antigen-agnostic enrichment of TRT from peripheral blood mononuclear cells (PBMC) by adopting a previously described protocol for antigen-reactive T cell generation using mature antigen-presenting cells (APC)^16,17^ (**Fig. 3A**). We established tumor PDO and a 2D line from one urothelial carcinoma patient and collected matched peripheral blood. Both tumor 2D lines and PDO expressed MHCI and MHCII (**Supplementary Fig. 4A-B**). Following, we cultured the PBMC in media containing APC differentiation and maturation proteins in the presence of IFN_γ_-treated PDO to increase antigen presentation and MHCI expression. PDOT co-culture enabled the generation of TRTs from peripheral blood, as evidenced by increased AIM expression upon PDO restimulation (**Supplementary Fig. 4C**). We generated ten T cell lines and compared their total T cell reactivity to the tumor 2D line, vehicle- or IFN_γ_-treated PDO (**Supplementary Fig. 4C-E**). Only one T cell line was reactive to the tumor 2D cells, which degranulated and produced GM-CSF and IFN_γ_ (**Supplementary Fig. 4F**). In contrast, 7/10 T cell lines showed reactivity to the vehicle- and IFN_γ_-treated tumor PDO (**Supplementary Fig. 4D-F**). TRT were distributed across CD8, DN and CD4 T cells (**Supplementary Fig. 4G**). These data suggest that tumor PDOs expand diverse repertoire of circulating TRTs with distinct reactivity for PDO and 2D tumor cells.

**Figure 3.**
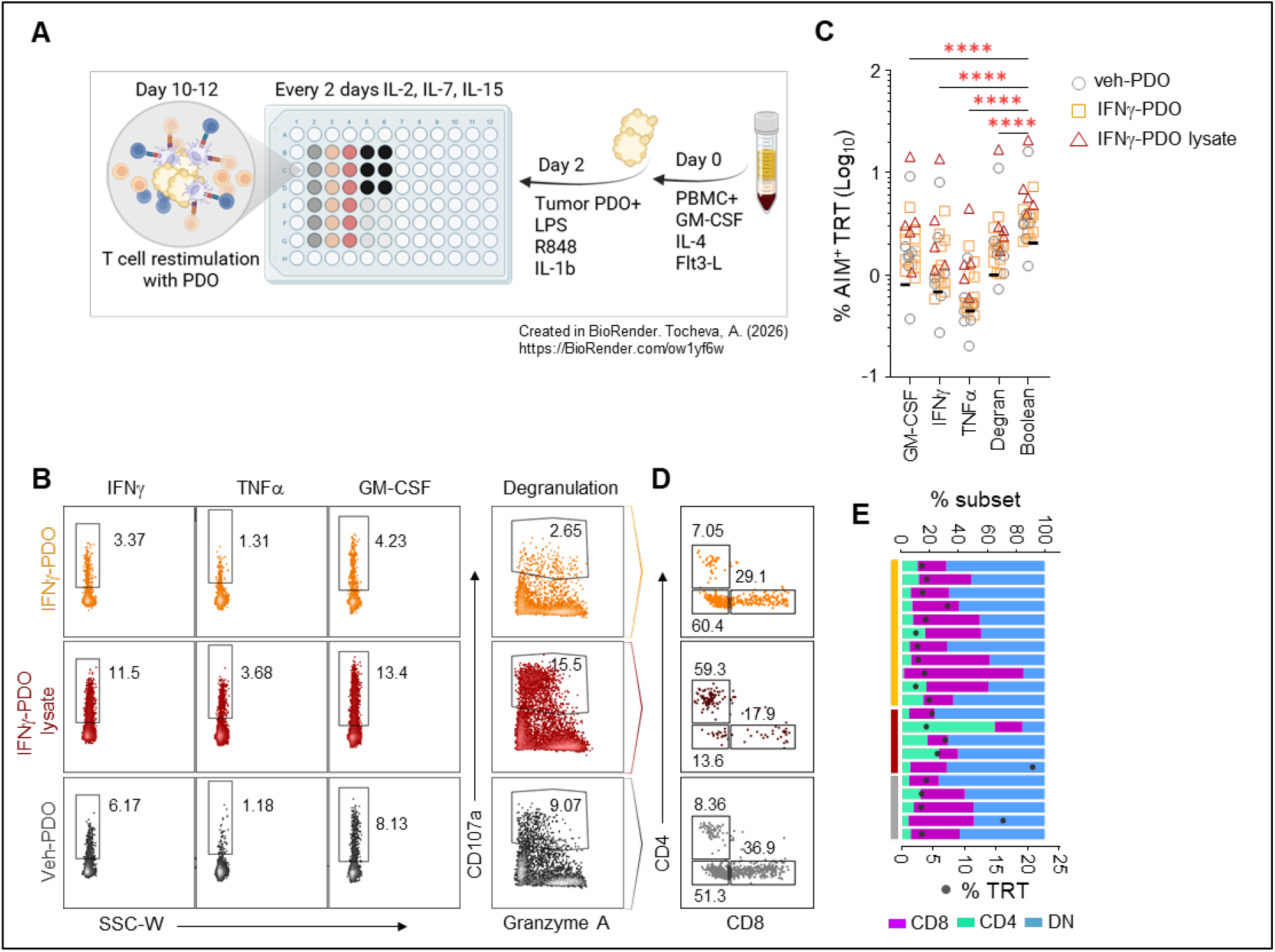
Enrichment of tumor PDO reactive T cells from peripheral blood. **A**, Experimental design for the generation of PDO-reactive T cells from peripheral blood. APC are differentiated from PBMC for 2 days followed by the addition of autologous tumor PDO and TLR ligands. Cells are fed every 2-3 days and the expansion of PDO reactive T cells is determined after day 10-12 by AIM flow cytometry. **B**, Representative flow cytometry plots of the expression of functional markers (y axis) in TRT generated from PBMC in response to IFN_γ_-treated PDO (orange), vehicle PDO (grey) or lysates prepared from IFN_γ_-treated PDO (red). **C**, Proportion of tumor-reactive T cells in TRT generated from peripheral blood identified by Boolean gating on all functional AIMs. Black lines correspond to the AIM average of unstimulated control cells. ****p≤0.0001 calculated using repeated measures one-way ANOVA with Dunnett correction for multiple comparisons. **D**, Representative plots of CD4, CD8 and DN T cell contribution to the blood-derived TRT repertoire in PDOTs identified by Boolean integration of all AIMs. **E**, CD4, CD8 and DN T cell distribution within TRT generated in response to PDO from peripheral blood.

IFN_γ_ stimulation increases antigen processing and presentation, and the expression of MHCI and MHCII^15^. Since all T cell lines in the first experiment were generated in response to IFN_γ_-treated PDO, we determined whether there are differences in the generation or functional repertoire of PBMC-derived TRT expanded in response to vehicle-treated PDO or IFN_γ_-treated PDO. Since PDO express high levels of PD-L1 upon IFN_γ_ stimulation, which may interfere with the generation of TRTs, we also used lysates from IFN_γ_-treated PDO. Following expansion, the PBMC-derived T cell lines were restimulated with their respective vehicle- or IFN_γ_-treated PDO, while lysate-reactive TRTs were restimulated with IFN_γ_-treated PDO (**Fig. 3B**). We detected TRTs in 11/12 lines in the IFN_γ_-PDO expanded cells, 5/5 lines in the lysate condition and 5/7 lines expanded in response to vehicle-treated PDO (**Fig. 3C**). Overall, TRT responses were consistently observed above unstimulated baseline across multiple PDO conditions, including vehicle-treated, IFN_γ_ -treated, and PDO lysate preparations (**Fig. 3C and Supplementary Fig. 4H**). Comparable levels of TRT were detected across conditions, indicating that TRT generation is robust and does not require IFN_γ_ conditioning or exogenous antigen loading. Analysis of TRT composition revealed that tumor-reactive cells were distributed across CD8, CD4, and DN T cells (**Fig. 3D-E**). Notably, a substantial fraction of TRT responses in the circulation was mediated by CD4 and DN T cells (**Supplementary Fig. 4I-J**). These findings indicate that peripheral blood contains a functionally diverse repertoire of tumor-reactive T cells that can be expanded using PDOT co-culture.

### PDOTs functionally stratify tumor-reactive T-cell responses to PD1 blockade and identify candidate combination checkpoint targets

Since PDOTs captured anti-tumor T cell reactivity across CD4, DN and CD8 T cells, we determined their utility to measure the functional TRT repertoire responding to PD1 blockade. PDOT co-cultures were performed in the presence of anti-PD1 antibody prior to intracellular AIM staining. Consistent with our previous findings, individual AIMs did not reliably identify PD1 blockade responsive TRT across PDOT (**Supplementary Fig. 5A**). Instead, Boolean integration of all AIMs identified anti-PD1 reactivity in 12/17 PDOTs, with three co-cultures exhibiting ≥25% increase in TRT responses (**Fig. 4A**). Notably, baseline tumor reactivity in the PDOTs was not a pre-requisite for PD1 sensitivity. We observed PDOT cultures with high baseline TRT activity that did not respond to PD1 blockade and PDOT cultures with absent or low T cell reactivity that showed increased TRT responses following treatment (**Fig. 4B**). In fact, PDOT responses to anti-PD1 (αPD1) were classified in four groups: (1) reactive non-responders (*i*.*e*. FGP315); (2) non-reactive responders (*i*.*e*. FGP341); (3) reactive responders (*i*.*e*. FGP269); and (4) non-reactive non-responders (*i*.*e*. FGP254) (**Supplementary Fig. 5B**). In the PDOT cultures exhibiting ≥25% reactivity (i.e. FGP269, FGP341 and FGP165), TRT were distributed across CD4, CD8 and DN T cells (**Fig. 4C-D**). Specifically, CD4 TRT were the major subset contributing to anti-PD1 response in FGP341 PDOTs, while CD8 and DN were the major responders in FGP269 and FGP165 (**Fig. 4D**). These results show that PDOT co-cultures enable functional stratification of tumor-reactive T-cell responses to PD1 blockade.

**Figure 4.**
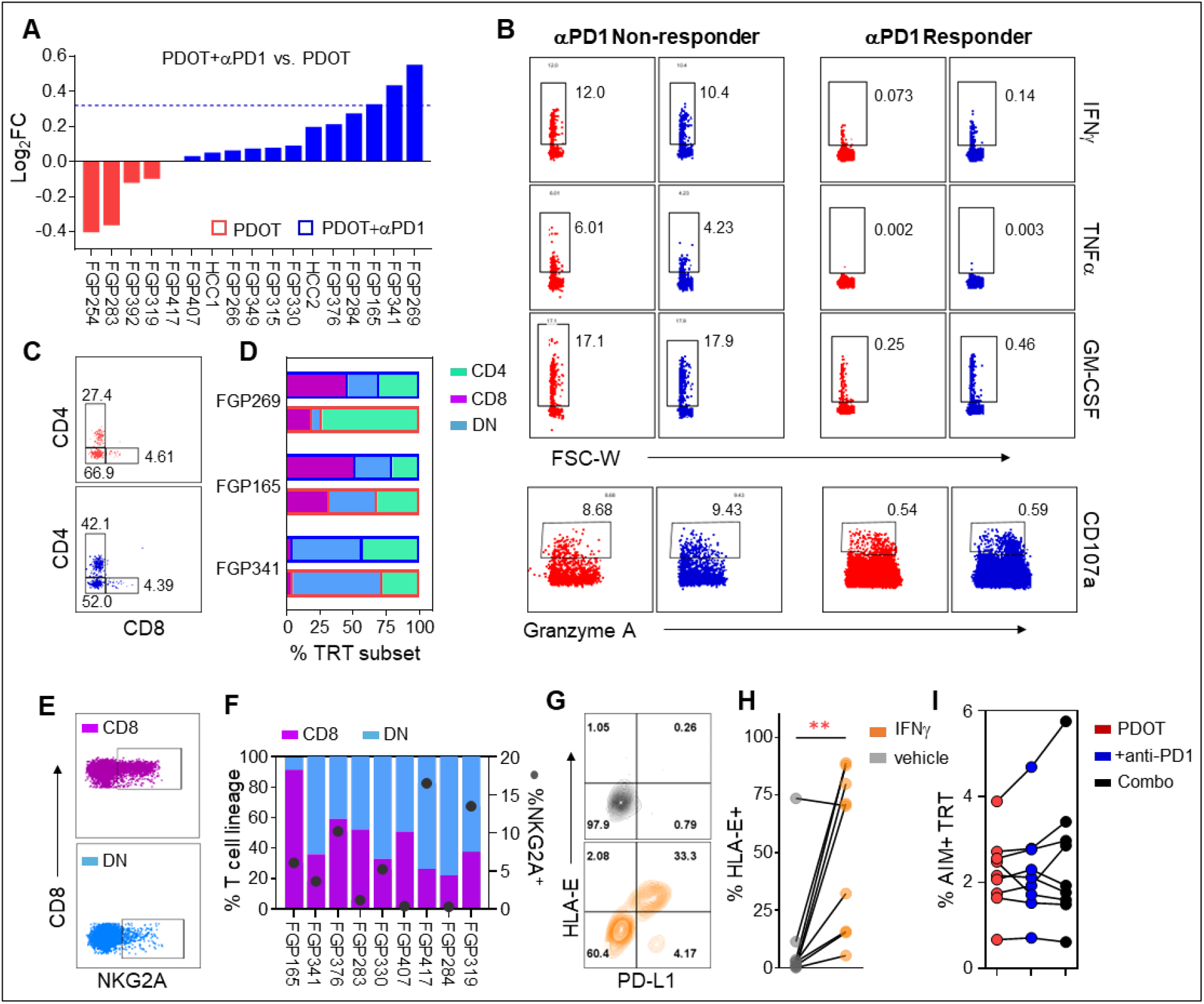
PDOT modeling of tumor-reactive T cell responses to immune checkpoint inhibitors. **A**, Magnitude of TRT responses in the PDOT in the absence (red bars) and presence (blue bars) of PD1 blocking antibodies. **B**, Representative flow cytometry plots of a PDOT with high baseline T cell reactivity that did not respond to PD1 blockade (left) and (right) PDOT with low baseline T cell reactivity that responded to PD1 blockade. **C**, Example flow cytometry plots showing CD4, CD8 and DN TRT distribution in response to PD1 blockade. **D**, Proportion of CD4, CD8 and DN TRT in PDOTs with ≥25% functional response to PD1 blockade. PDOT (bars outlined in red) and PDOT+aPD1 (bars outlined in blue). **E**, Example flow cytometry plots of NKG2A expression in CD8 and DN T cells from TIL. **F**, Proportion of CD8 and DN T cells within TIL that express NKG2A (left y axis) and raw percent of NKG2A positive T cells (right y axis). **G**, Example flow cytometry pots of HLA-E expression in the PDO in response to vehicle (grey) or IFNg (orange). **H**, Quantified expression of HLA-E from G across multiple PDO; **p<0.01 using paired Student’s t test. **I**, Proportion of TRT in response to PDOT (red), PDOT+αPD1 (blue) or PDOT+αPD1/αNKG2A (black) (n=9).

We next asked whether PDOTs could identify additional inhibitory pathways that limit TRT responses. We noticed that NKG2A was highly expressed on CD8 and DN T cells in TIL cultures (**Fig. 4E-F**) and its ligand HLA-E was upregulated by IFN_γ_ stimulation in the PDO (**Fig. 4G-H**). Therefore, we determined whether combined blockade of PD1 and NKG2A will increase T cell responses in PDOTs. Out of nine tested PDOT co-cultures, the combination treatment increased TRT responses in four PDOTs generated from patients with urothelial (n=1), lung (n=2) and colorectal cancers (n=1) (**Fig. 4I** and **Supplementary Fig. 5C**). Collectively, these results demonstrate that PDOTs can be leveraged to functionally stratify anti-tumor T cell responses to PD1 blockade and to identify patient-specific combination checkpoint strategies that enhance anti-tumor T cell activity.

## Discussion

Our study establishes PDOT co-cultures as a pre-clinical platform to assess the magnitude and functional diversity of tumor-reactive T cell responses. While PDOT co-cultures have been previously used to identify TRTs, they have primarily focused on CD8 T cell reactivity^3,4^. In contrast, the contribution of DN and CD4 T cells remain under-explored^18-21^, and preclinical models that evaluate their functions are limited. Moreover, the use of single activation marker readouts has limited the detection of the full TRT repertoire and obscured the functional heterogeneity of immune checkpoint inhibitor responses. Here, we show that multiplex immune profiling of PDOT co-cultures is necessary to capture the breadth and functional diversity of tumor-reactive T cells.

Analytically, we demonstrate that individual activation markers underestimate TRT responses. Prior studies have relied on IFN_γ_, CD107a or CD137 to identify TRTs. However, these single readouts were non-overlapping across reactive T cells in the PDOTs. Boolean integration of multiple AIMs identified significantly more TRTs compared to any individual marker. This likely reflects both functional exhaustion from chronic antigen exposure and prolonged *in vitro* culture, as well as intrinsic differences in effector programs across CD4, CD8 and DN T cells. These findings highlight the need for multiplex functional approaches to accurately define the TRT repertoire.

Our work further highlights that in addition to CD8s, PDOTs enable the identification of CD4 and DN TRTs in both tumor infiltrating lymphocytes and peripheral blood. Tumor PDO expressed both MHCI and MHCII molecules, and MHCI blocking antibody abrogated CD8 and DN T cells responses while preserving CD4 reactivity, supporting antigen-dependent recognition across T cell lineages. Notably, PDOTs do not contain professional antigen presenting cells suggesting that tumor PDO can directly present antigens to CD4 T cells in this system. Meanwhile, DN T cells have strong cytotoxic functions and were among the highest cytokine producing population. While both _γ_δ and αβ T cells contribute to the DN compartment, the MHCI restriction of DN TRT observed here suggests a predominant αβT cell origin. Furthermore, PDOTs generated and identified more peripheral blood TRTs compared to autologous 2D cells. Since PDOs better preserve the genomic diversity of the primary tumor than 2D lines, this likely reflects increased antigen diversity that supports a broader TRT repertoire, although further studies are needed to validate this. Collectively, these findings establish PDOTs as a platform to identify functional TRT across T cell lineages.

Translationally, the PDOT platform has several applications. PDOTs enable functional interrogation of tumor-T cell interactions and responses to immunotherapy. We show that PDOTs capture heterogeneous TRT responses to PD1 blockade and that baseline tumor reactivity was not a pre-requisite for sensitivity to PD1 immune checkpoint inhibitor treatment. Both reactive and non-reactive PDOT cultures exhibited variable reactivity to PD1 blockade. Additionally, PDOTs supported the identification of candidate combination immune checkpoint inhibitors that improved TRT reactivity. We observed increased NKG2A expression in TILs and HLA-E upregulation in the tumor PDO and combined NKG2A and PD1 blockade improved TRT responses in a subset of PDOT cultures. While further studies are required to determine the accuracy of PDOT models to predict *in vivo* patient responses, these findings demonstrate that PDOTs can be leveraged to functionally stratify tumor-T cell responses and to prioritize patient-specific immunotherapeutic strategies.

## Limitations

Our study has limitations. While PDOT co-cultures provide a tractable platform to study tumor-T cell interactions, they represent a reductionist system that does not fully recapitulate the complexity of the tumor microenvironment. Specifically, PDOTs lack stromal and myeloid components that can influence antigen presentation, T cell activation and responses to ICIs. Incorporating stromal and myeloid components will significantly improve current PDOT platforms. Although we observed CD4 T cell responses in the PDOT in the absence of professional antigen-presenting cells, suggesting that tumor cells can directly present antigen via MHCII, the relevance of this interaction warrants further investigation. Additionally, T cells in the PDOT are subject to *in vitro* expansion, which may change their functional state and immune checkpoint expression profiles compared to their *in vivo* counterparts. TRT reactivity in this study was based on the functional analysis of activation induced markers and we did not directly assess tumor cell killing in PDOT co-cultures or following immunotherapy treatment. Whether functional reactivity translates into effective tumor killing, especially in response to ICI treatment, requires further validation. Finally, although our findings support the identification of candidate combination checkpoint strategies such as PD1 and NKG2A blockade, these observations require validation in larger cohorts. Despite these limitations, PDOTs provide a controlled and scalable platform to functionally interrogate tumor-reactive T cell responses, identify tumor-intrinsic mechanisms that perturb T cell mediated anti-tumor immunity and prioritize patient-specific immunotherapeutic strategies.

## Methods

### Tumor organoid generation and immunophenotyping

Patient-derived tumor tissue (∼0.25 – 1 cm^3^), which was sourced from consenting patients from the Mount Sinai Biorepository under Institutional Review Board (IRB) protocol, was minced to 1-2 mm^3^ pieces and digested in gentleMACS dissociator using the MACS Tumor Dissociation Kit (Miltenyi, 130-095-929) according to manufactures instructions. Ice cold RPMI was added to the digest and samples were passed through a 70 μm nylon cell strainer and centrifuged for 5 min at 350g. Pellets with erythrocyte contamination were treated with ACK lysis buffer, washed with ice cold RPMI and centrifuged. The single cell dissociate was resuspended in cold tissue-specific organoid media as described previously^13,22^, then mixed with Matrigel (Corning 354230) at a ratio of 1:2 for a final concentration of 1 x 10^6^ cells/mL. The cell/Matrigel suspension was added to pre-warmed tissue culture plates and the plates were inverted at 37°C, 5% CO_2_ humidified incubator. After 30 min, pre-warmed tissue-specific organoid media was added to each well. Medium was changed twice a week and confluent cultures were passaged for expansion. For PDO cryopreservation, the PDOs were first recovered from the Matrigel using Cell Recovery Solution (Corning) and then preserved in Bambanker freezing media (Bulldog-Bio). Phase-contrast light microscopy of organoid cultures was carried out using an EVOS XL CORE Imaging System (Invitrogen). For immunophenotyping, tumor PDO were recovered from Matrigel with ice-cold cell recovery solution and dissociated to single cell suspension with TrypLE at 37C for 1min. For the staining, 3 x 10^5^ cells were transferred to 12 x 75mm polystyrene tubes, washed once with PBS and resuspended in 70 μL Zombie NIR (BioLegend) at 1:3000 final dilution in PBS. The cells were incubated for 15 min at room temp in the dark, then washed with FACS staining buffer (2% FBS (v/v), 2mM EDTA in PBS) and spun for 5 min at 350g. Pellets were resuspended with 50 μl 1:50 Human TruStain FcX (Biolegend) prepared in FACS staining buffer and incubated for 20 min on ice in the dark. Without washing, 50 μL fluorochrome conjugated anti-human antibody mix containing anti-MHCI-PECy5, anti-PD-L1-PE-Dazzle 594, anti-MHCII-PECy7 and anti-human HLA-E-PE (Biolegend) was added to each tube and incubated for 30 min on ice in the dark. The samples were washed twice and fixed with 1% PFA (v/v) in PBS prior to acquisition with Cytek Aurora 5-laser spectral cytometer (Cytek Biosciences). Flow cytometry data cleanup was performed with FlowAI and downstream analysis was performed using FlowJo v10 software (BD Biosciences).

### Organoid Media recipes

#### Base media

Advanced DMEM (ThermoFisher Scientific, Catalog #12634-034), Noggin/ R-Spondin (from ATCC# CRL-3276, LWRN cell media), B27 supplement (ThermoFisher Scientific, Catalog #17504-044), Glutamax-I (ThermoFisher Scientific, Catalog #35050-061), Hepes (ThermoFisher Scientific, Catalog # 15630-080), Primocin (Inivogen, Catalog # ant-pm-1) and penicillin/streptomycin (ThermoFisher Scientific, Catalog #15140-122).

#### Lung media

Base media with Recombinant Human FGF-7 ([25ng/mL], PeproTech, Catalog # 100-19), Recombinant Human FGF-10 ([20ng/mL], PeproTech, Catalog #100-26), SB202190 ([10μM MilliporeSigma, Catalog #S7067), Y-27632 ([10μM], Selleck Chemicals, Catalog #S1049), A-83-01 ([500nM], Tocris, Catalog #2939), Nicotinamide, ([10mM], MilliporeSigma, Catalog # N0636), N-acytlecysteine ([1.25mM], MilliporeSigma, Catalog #A9165).

#### Colon media

Base media with Recombinant Human FGF-basic ([1ng/mL], PeproTech, Catalog #100-18B), Recombinant Human FGF-10 ([20ng/mL], PeproTech, Catalog #100-26), PGE2 ([1 μM], R&D Systems, Catalog # 2296/10), SB202190 ([10μM MilliporeSigma, Catalog #S7067), Recombinant Mouse EGF ([50ng/mL], ThermoFisher Scientific, Catalog#PMG8043), Y-27632 ([10μM], Selleck Chemicals, Catalog #S1049), A-83-01 ([500nM], Tocris, Catalog #2939), Nicotinamide, ([10mM], MilliporeSigma, Catalog # N0636),N-acytlecysteine ([1.25mM], MilliporeSigma, Catalog #A9165), NRG ([10ng/mL], PeproTech, Catalog #100-03), Gastrin ([10nM], MilliporeSigma, Catalog # 05-23-2301).

#### Testicular media

Base media with Recombinant Human FGF-basic ([1ng/mL], PeproTech, Catalog #100-18B), Recombinant Human FGF-10 ([20ng/mL], PeproTech, Catalog #100-26), PGE2 ([1 μM], R&D Systems, Catalog # 2296/10), SB202190 ([10μM MilliporeSigma, Catalog #S7067), Recombinant Mouse EGF ([50ng/mL], ThermoFisher Scientific, Catalog#PMG8043), Y-27632 ([10μM], Selleck Chemicals, Catalog #S1049), A-83-01 ([500nM], Tocris, Catalog #2939), Nicotinamide, ([10mM], MilliporeSigma, Catalog # N0636), N-acytlecysteine ([1.25mM], MilliporeSigma, Catalog #A9165), NRG ([10ng/mL], PeproTech, Catalog #100-03), Follicle stimulating hormone ([20 mIU/mL-1], Abcam, Catalog #ab51888), human chorionic gonadotropin ([5 IU/L], MilliporeSigma, Catalog #C1063).

#### Kidney media

Base media with Recombinant Human FGF-basic ([1ng/mL], PeproTech, Catalog #100-18B), Recombinant Human FGF-10 ([20ng/mL], PeproTech, Catalog #100-26), PGE2 ([1 μM], R&D Systems, Catalog # 2296/10), SB202190 ([10μM MilliporeSigma, Catalog #S7067), Recombinant Mouse EGF ([50ng/mL], ThermoFisher Scientific, Catalog#PMG8043), Nicotinamide, ([10mM], MilliporeSigma, Catalog # N0636),N-acytlecysteine ([1.25mM], MilliporeSigma, Catalog #A9165), Y-27632 ([10μM], Selleck Chemicals, Catalog #S1049), A-83-01 ([500nM], Tocris, Catalog #2939), NRG ([10ng/mL], PeproTech, Catalog #100-03.

#### Liver media^23^

Base media with N2 supplement (ThermoFisher Scientific, Catalog #17502048), N-acytlecysteine ([1.25mM], MilliporeSigma, Catalog #A9165), Nicotinamide, ([10mM], MilliporeSigma, Catalog # N0636), Recombinant Human [Leu15]-Gastrin I ([10nM], MilliporeSigma, Catalog #G9145), Recombinant Human EGF ([50ng/ml], ThermoFisher Scientific, Catalog #PHG0311), Recombinant Human FGF-10 ([20ng/mL], PeproTech, Catalog #100-26), Recombinant Human HGF ([25ng/ml], ThermoFisher Scientific, Catalog #100-39H), Forskolin ([10μM] ThermoFisher Scientific, Catalog #J63292.MA), A-83-01 ([5μM], Tocris, Catalog #2939), Recombinant Human Noggin ([25ng/ml], ThermoFisher Scientific, Catalog #120-10C), Y-27632 ([10μM], Selleck Chemicals, Catalog #S1049), R-Spondin1 (from MilliporeSigma, Catalog #SCC11 or R-Spondin1 293T cell conditioned media), Wnt (from ATCC# CRL-2647, L Wnt-3A cell conditioned media). L Wnt-3A, Noggin and Y-27632 were removed from media after initial 3 weeks of culture.

### TIL generation

Paired TILs were generated by culturing 1 million cells from the tumor dissociate at 37°C, 5% CO_2_ in TIL media containing RPMI supplemented with 10% (v/v) human AB serum, penicillin/streptomycin, non-essential ammino acids, sodium pyruvate and 6000IU/mL IL-2 (Aldesleukin, provided by Clinigen). Fresh TIL media was replenished three times/week. Seven days after culture initiation, IL-2 concentration was reduced to 3000IU/ml and T cell expansion was augmented with 25 μL/well of ImmunoCult Human CD3/CD28/CD2 T Cell Activator (STEMCELL Technologies, 10970). Cultures were passaged when media color changes and TILs were cryopreserved at the end of week 3 in 90% human AB serum, 10% DMSO (v/v) in liquid nitrogen.

### Generation of tumor 2D and PDO-reactive PBMC-T cells

Fresh peripheral blood was kept at room temperature and PBMC were isolated within 4 hours of collection with the Lymphoprep Density Gradient Medium (STEMCELL Technologies). PBMC were either used fresh or frozen in 90% human serum, 10% DMSO (v/v) and kept in liquid nitrogen for long term storage. Frozen PBMC were recovered in pre-warmed TIL media without cytokines with 1 μg/mL DNase I and centrifuged for 10 min at 400g. To generate tumor 2D and PDO-reactive PBMC-T cells, pelleted PBMC were resuspended in warm X-VIVO 15 media (Lonza Bioscience) at 10^6^ cells/mL and 100 μL (10^5^ cells) of the cell suspension was added to each well of a 96-well U bottom plate. The 2X cytokine media was prepared with X-VIVO15 containing 2000 IU/mL GM-CSF, 1000 IU/mL IL-4, 100 ng/mL Flt3-L as previously described, and 100 μL were pipetted to each PBMC well. The cells were cultured for 24h in a humidified incubator at 37°C with 5% CO_2_ followed by 100 μL of media replacement from each well with 2X adjuvant solution consisting of X-VIVO15 with 20 μM R848, 200 ng/mL LPS, 20 ng/mL IL-1β and either 1μM vaccine CEFT peptide pool from JPT and either 2D tumor cell lines, tumor PDO, or tumor PDO lysates. 2D tumor cells and 3D tumor organoids were pre-stimulated with 200 ng/mL IFNγ for 24 h where indicated and used at 2 x 10^4^ single cell equivalents per well. Tumor organoid lysates were prepared by sonication of organoids in PBS and used at a final concentration of 5 μg/mL. The cells were cultured for 24h in a humidified incubator at 37°C with 5% CO_2_ prior to media replacement with 100 μl/well of 2X feeding media consisting of RPMI supplemented with 10% (v/v) human AB serum, penicillin/streptomycin, non-essential amino acids, sodium pyruvate, 20 IU/mL IL-2 (Aldesleukin, provided by Clinigen), 20 ng/mL IL-7 (Peprotech) and 20 ng/mL IL-15 (Peprotech). The cells were cultured in a humidified incubator at 37°C with 5% CO_2_. PBMC-T cell lines were re-stimulated each week using the corresponding (peptide), 2D tumor cell line, tumor PDO, or PDO lysate. Media replacement was performed every 2-3 day. Reactivity of PBMC-T lines was assessed in PDOT co-culture at D10. PBMC-T were frozen in 90% human serum, 10% DMSO (v/v) and kept in liquid nitrogen for long-term storage.

### Whole exome sequencing analysis

Paired-end FASTQ files were aligned to the human reference genome (hg38) using BWA-MEM^24^, followed by sorting and duplicate marking. Somatic variants were called using GATK Mutect2^25^ in tumor-only mode with a panel of normals (PON) and a germline resource (gnomAD)^26^, and filtered using FilterMutectCalls. Only variants passing all filters were retained. Variants were restricted to chromosomes 1–22 and X, intersected with exome capture target regions, and filtered to retain high-confidence somatic single nucleotide variants (SNVs) with sequencing depth ≥10. Variant allele frequencies at a pooled set of somatic SNV sites generated across samples were extracted for each sample, and pairwise Pearson correlation coefficients were computed. The correlation matrix was visualized as a heatmap following hierarchical clustering with the Ward linkage method. Variant data preprocessing was performed in Python, Pearson correlation coefficients were computed using NGSCheckMate^27^, and visualization was performed in R.

### PDOT co-cultures and spectral cytometry

Expanded TIL and PBMC-T lines were rested in cytokine-free TIL media for 24h prior to co-culture with paired PDO. PDO were stimulated with 200 ng/mL IFNγ for 24h prior to co-culture to enhance antigen presentation and PD-L1 expression. PDO recovery from the Matrigel culture was performed as described above aiming to preserve 3D structure in the co-cultures. Washed PDO were resuspended in warm TIL media without cytokines at 5 x 10^5^ single cell equivalents per mL. For MHCI blocking experiments, mouse-anti human HLA-A,B,C (clone W6/32, Biolegend) was included at 10 μg/mL final concentration for 30min at 37°C with 5% CO_2_ prior to co-culture with TIL. For T cell preparation, the cells were washed with warm TIL media for 4 min at 400g and resuspended at 1 x 10^6^ cells/mL in warm TIL media with 2 μg/mL mouse anti-human CD28 (Biolegend). The T cell mixture was transferred to 5 mL FACS tubes at 300 μL/tube. For PD1 blockade, T cells were incubated for 30min at 37°C with 5% CO_2_ with 20μg/mL anti-PD1 blocking antibody (Nivolumab). PDO were combined with the T cells by adding 100 μL of PDO suspension to each T cell tube. BV785 mouse anti-human CD107a (Biolegend) was diluted 1:50 and added at 50 μL/tube. PDOT cell co-culture tubes were centrifgued for 15 s at 200g to initiate PDO-T cell contact and placed in a humidified incubator at 37°C with 5% CO_2_ for 1 h. Brefeldin and Monensin (both from Biolegend) were added to each tube at a final concentration of 0.5X each, and the volume was adjusted to 500 μL/tube. PDOT co-culture tubes were placed in a humidified incubator at 37°C with 5% CO2 for 15h prior to processing for intracellular cytokine staining (ICS). For the ICS, PDOT co-cultures were washed with PBS for 4 min at 400g. The pellets were resuspended with pre-warmed TrypLE (Gibco) at 200 μL/tube and incubated for 1 min at 37°C. The cells were washed with 3 mL PBS at room temperature and the pellets were resuspended with 70 μL 1:3000 Zombie NIR (Biolegend) prepared in PBS. The PDOT tubes were incubated 15 min at room temp in the dark, then washed with FACS staining buffer (2% FBS (v/v), 2mM EDTA in PBS) and spun at 400g for 4 min. The pelleted cells were resuspended with 50 μl 1:50 Human TruStain FcX (Biolegend) in FACS staining buffer and incubated for 20 min on ice in the dark. Without washing, 50 μL of antibody staining mix containing BUV395 anti-CD45, BUV496 anti-CD3, BV570 anti-CD4, Pacific Blue anti-CD8, BUV805 anti-CD56, BV711 anti-PD1, FITC anti-LAG3, BV421 anti-2B4, PE-Cy5 anti NKG2A, PerCP-Cy5.5 anti-TIM3, APC anti-TIGIT, and PE-Cy7 anti-GITR was added to each tube and incubated for 30 min on ice in the dark. The samples were washed twice with FACS staining buffer for 4 min at 400g and the pellets were resuspended in 250μl Fixation Buffer (Biolegend) and incubated for 20 min at room temperature in the dark. Tubes were washed with 1X Permeabilization Wash Buffer (Biolegend) for a total of 3 washes. The pellets were resuspended in 50μl of 1X Permeabilization Wash Buffer (Biolegend) containing Pacific Blue anti-CD8, AlexaFluor-700 anti-granzyme A, BUV737 anti-IFNγ, BV605 anti-TNFα, PE anti-CD137 and PE-Dazzle594 anti-GM-CSF. Following the stain, 1X Permeabilization Wash Buffer (Biolegend) was added to each tube and the tubes were spun for 4 min at 400g. The pellets were washed in FACS acquisition buffer (0.2% FBS (v/v), 2mM EDTA in PBS), resuspended in 100μL FACS acquisition buffer and acquired using a 5-laser Cytek Aurora spectral cytometer (Cytek Biosciences). Data were analyzed using FlowJo v10 software (BD Biosciences). Statistical analyses were carried out using GraphPad Prism v10.1.0.

## Data availability

All data supporting the findings of this study are available within the paper and its Supplementary Information, and available upon request from the corresponding authors.

## Acknowledgements

We sincerely thank the patients for their invaluable contribution to our research and the Dean’s Biorepository core at the Icahn School of Medicine at Mount Sinai. Research reported in this publication was supported by the Flow Cytometry CoRE (RRID:SCR_027701) at the Icahn School of Medicine at Mount Sinai and by the National Cancer Institute of the National Institutes of Health under award number P30 CA196521. The content is solely the responsibility of the authors and does not necessarily represent the official views of the National Institutes of Health.

## Research Funding

This work was supported by the Department of Genetics and Genomic Sciences and CIC Project Pilot award to AST and BH, National Cancer Institute NCI (NCI, R00-CA230384) to BH, in part by the Mount Sinai Tisch Cancer Center and ConduITS developmental Fund Award to DS and AST, and in part by the American Lung Association Susan Rappaport Lung Cancer Discovery Award to AMT. DS was supported by the Department of Defense (RA220126), National Cancer Institute (NCI, R01-CA285580-01) and RSG-23-1151048-01-IBCD from the American Cancer Society [https://doi.org/10.53354/ACS.RSG-23-1151048-01-IBCD.pc.gr.175471]. DS and AST are supported by the National Cancer Institute (NCI, R01-CA285425).

## Author contributions

Conceptualization: BH and AST; Experimental design: EM, JCF, BH and AST; Methodology/Investigation: EM, JCF, SN, GLG, EVA, RS and SS; Data analysis: EM, EA, WL, AMT and AST; Sample procurement: LZ, FRH, JPS, KB, RB, AH, JCF and BH; Supervision/Administration and funding: BH and AST; Data interpretation: MW, AH, DS, AMT, BH and AST; Writing: EM, BH and AST. All authors reviewed and approved the final manuscript.

