## Supplemental Figures for "Organoid-T cell co-cultures functionally stratify tumor-reactive T cells and their responses to immune checkpoint inhibitors"

^3^ Current address: Englander Institute for Precision Medicine, Weill Cornell Medicine, New York, NY, USA; Department of Physiology and Biophysics, Weill Cornell Medicine, New York, NY, USA.

*These authors contributed equally

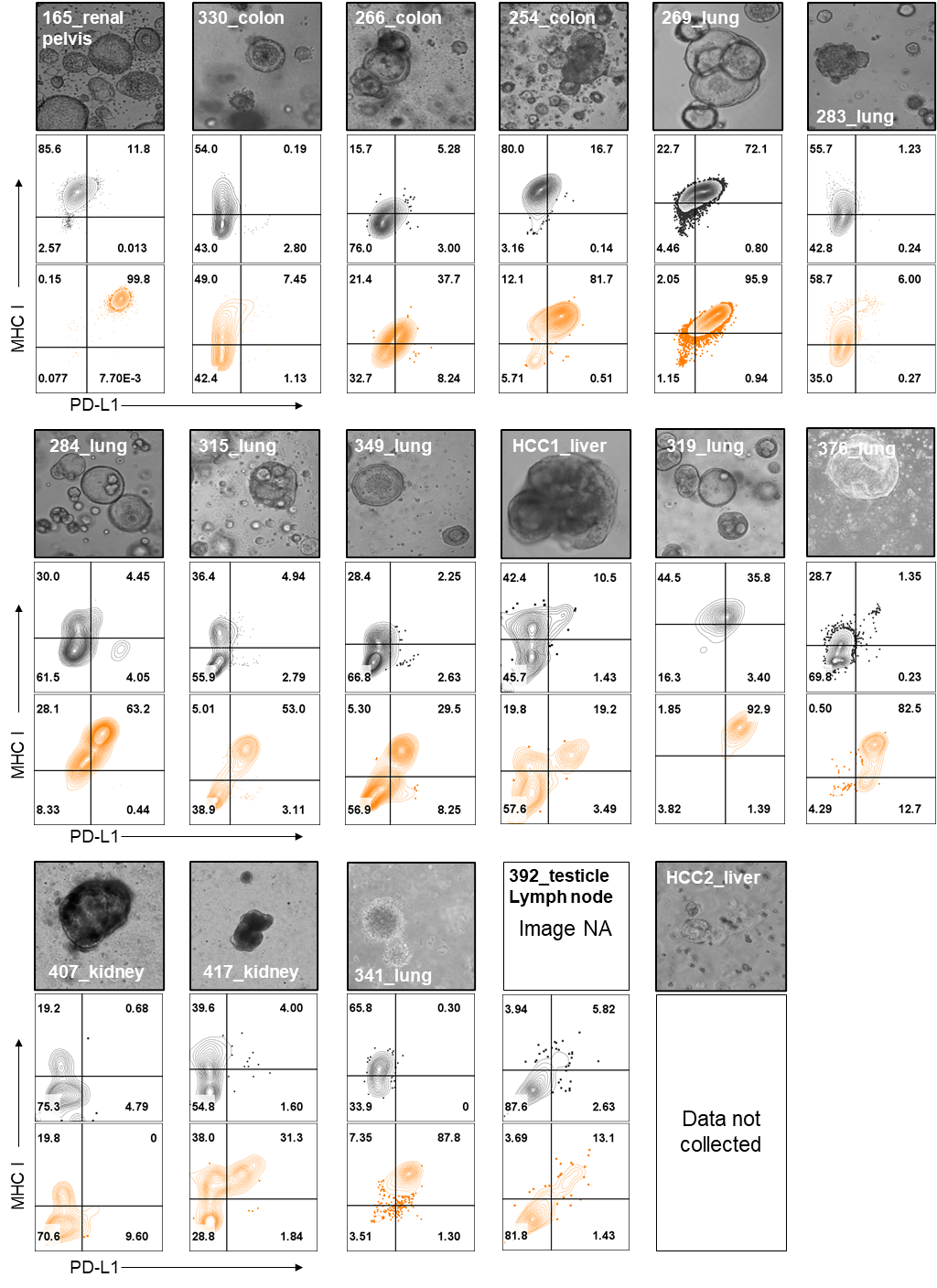

**Supplementary Figure 1. Brightfield micrographs of tumor PDO and PDO flow cytometry plots of PD-L1 and MHCI expression.**

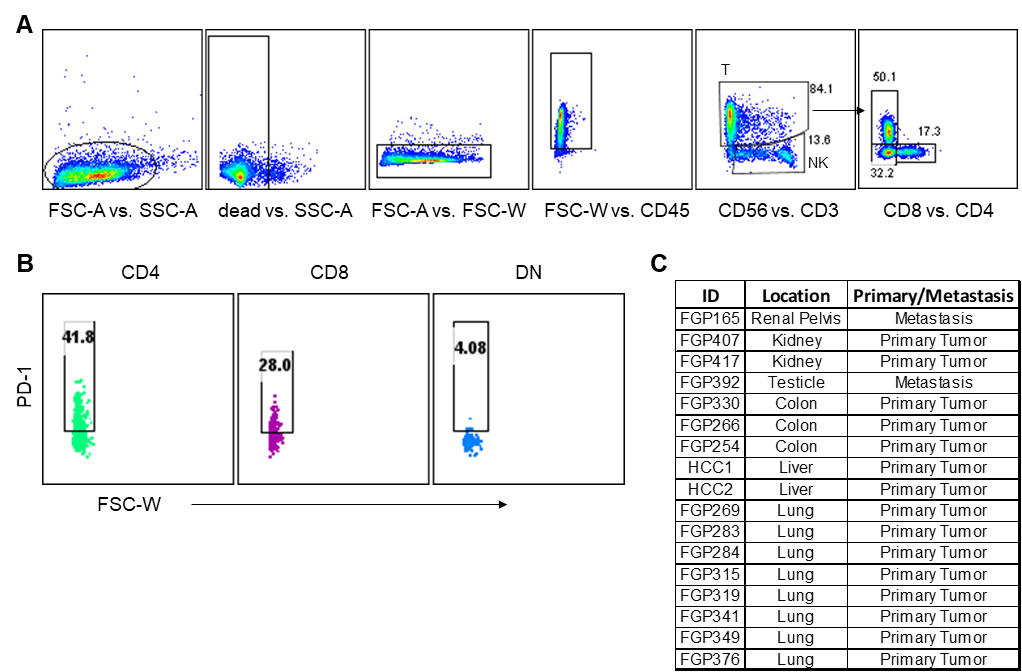

**Supplementary Figure 2. Gating strategy for TIL immunophenotyping**. **A,** TIL gating strategy to quantify NK cells and CD4, CD8, DN T cells. **B,** Representative plots of PD1 expression (%) in CD4, CD8 and DN T cells. **C,** Demographic characteristics of the patients.

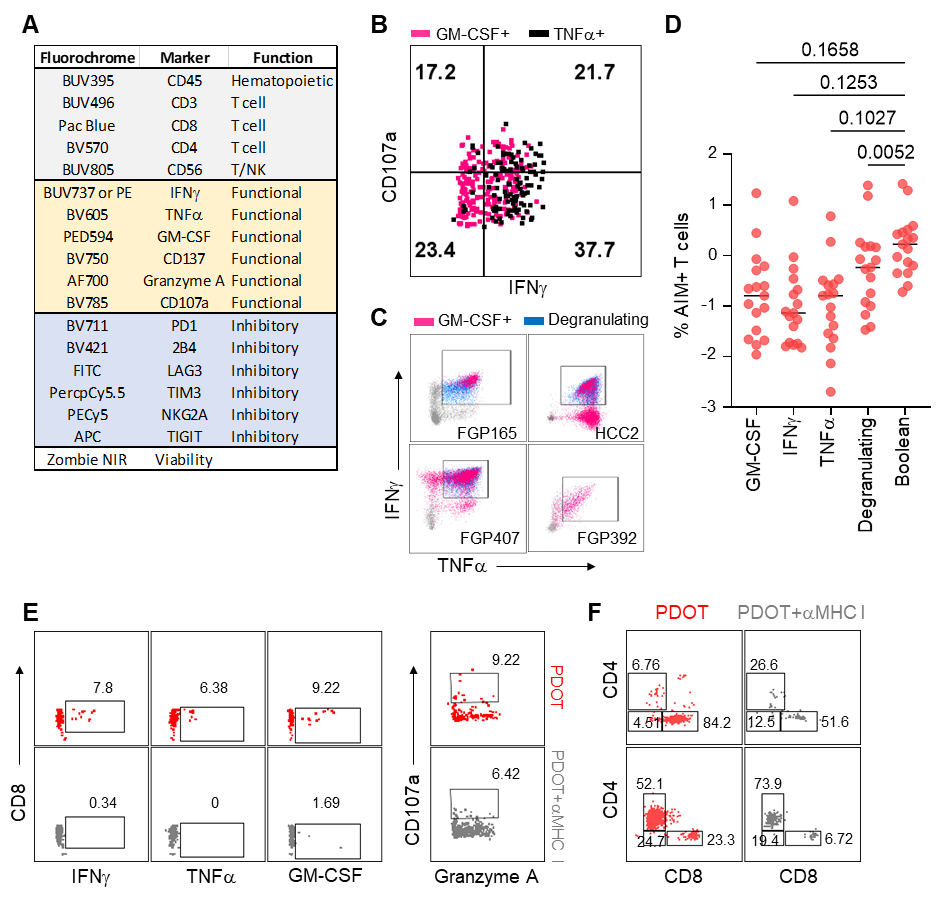

**Supplementary Figure 3. Activation induced markers for immunophenotyping of tumor-reactive T cells in the PDOT co-cultures**. **A,** Functional TIL panel. **B,** Representative dot plots of functional marker expression in TRT from PDOT of FGP315 or **C,** from T cells upon PMA/Ionomycin stimulation showing the exclusive and combined production of each AIM. **D,** % AIM+ T cells in PMA/Ionomycin-stimulated TIL; p-values calculated using repeated measures one-way ANOVA with Dunnett correction for multiple comparisons. **E,** Representative flow cytometry plots of DN TRT in the PDOT (red) and in response to anti-MHCI blocking antibody (grey) showing that DN TRT responses are MHCI-dependent. **F,** Representative flow cytometry plots gated on total AIM+ TRT showing reduced proportion of CD8 TRT upon MHCI blockade but not CD4 TRT.

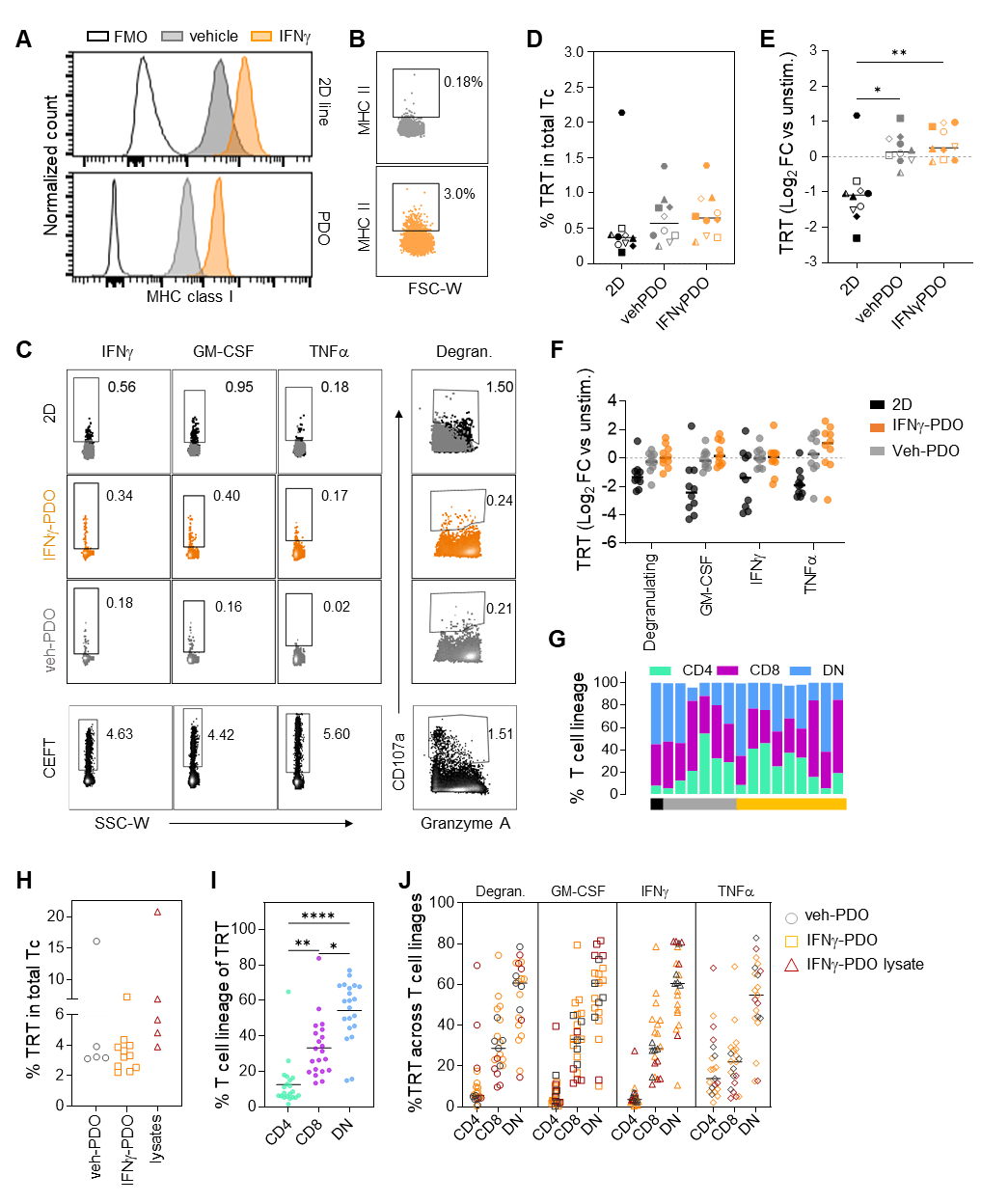

**Supplementary Figure 4. Generation of tumor PDO-reactive T cells from peripheral blood. A,** MHC class I expression in tumor PDO or matched 2D line following overnight stimulation with vehicle (grey) or IFNγ (orange). **B,** Flow cytometry plots of MHCII expression in the PDO used to generated blood-derived TRTs. C, Representative flow cytometry plots of AIM+ blood-derived TRT detected in response to tumor 2D lines stimulation, vehicle- or IFNγ-treated PDO. CEFT pool was used as a positive control. **D,** Proportion and (**E**) Log_2_FC relative to unstimulated cells of blood TRT detected in response to tumor 2D line, vehicle- or IFNγ-treated PDO across 10 blood-generated T cell lines. *p≤0.05, **p≤0.01 calculated using repeated measures one-way ANOVA with Tukey correction for multiple comparisons. **F,** Individual AIM expression in blood-derived TRT lines in response to tumor 2D lines stimulation, vehicle- or IFNg-treated PDO. **G,** Proportion of T cell lineage distribution of the TRT identified from blood-derived T cell lines in response to 2D lines stimulation, vehicle- or IFNγ-treated PDO. **H,** Proportion of blood derived TRT generated in response to vehicle-PDO (grey), IFNγ-treated PDO (orange) or lysates from IFNγ-treated PDO (burgundy). **I,** Proportion of CD4, DN and CD8 T cells in TRT from H. *p≤0.05, **p≤0.01 and ****p≤0.0001 calculated using repeated measures one-way ANOVA with Tukey correction for multiple comparisons. **J,** Proportion of CD4, DN and CD8 T cells in TRT contributing to individual AIM expression.

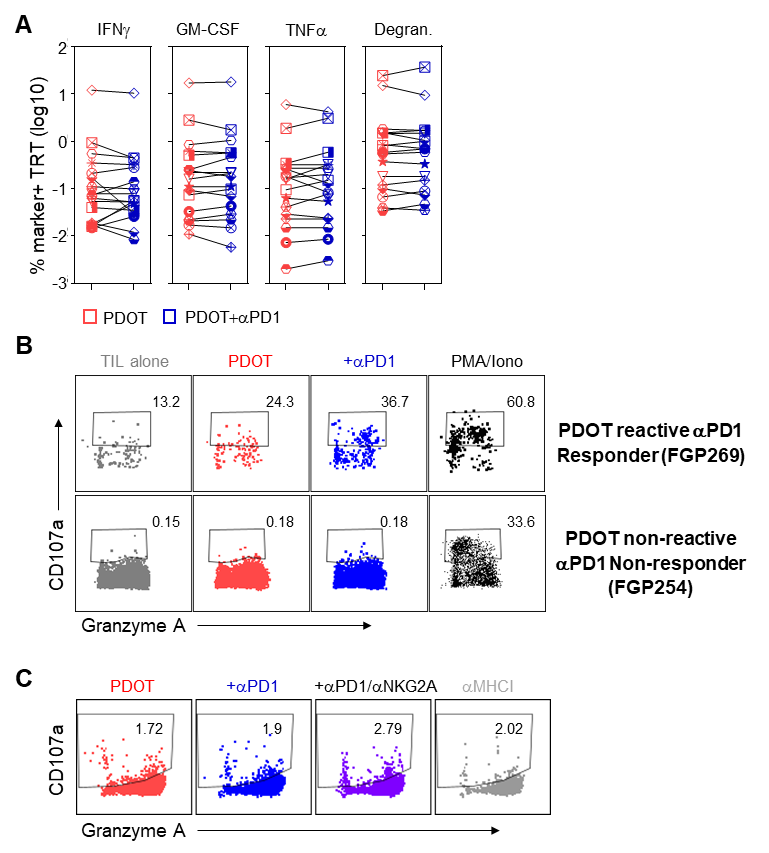

**Supplementary Figure 5. PDOT responses to immune checkpoint inhibitors.** **A,** Proportion of individual AIM+ TRT in the PDOT or PDOT treated with anti-PD1 antibody. **B,** Representative flow cytometry plots of PDOT with baseline tumor reactivity that responded to PD1 blockade by increasing AIM expression (top) and (bottom) PDOT lacking baseline tumor reactivity that did not respond to PD1 blockade. **C,** Representative flow cytometry plots of PDOT co-culture showing that the combined blockade of PD1 and NKG2A improves TRT responses in MHCI-dependent manner.
